# More isn’t always better: Too much exoskeleton torque can disrupt balance

**DOI:** 10.64898/2026.06.02.729541

**Authors:** Joon Han Kim, Rish Rastogi, Giovanni Martino, Owen N. Beck, Max K. Shepherd, Gregory S. Sawicki, Lena H. Ting, Kristen L. Jakubowski

**Affiliations:** College of Arts and Sciences, Emory University, Atlanta, GA, USA; Department of Physical Therapy and Rehabilitation Science, Northeastern University, Boston, MA, USA; Department of Biomedical Sciences, University of Padova, Padua, Italy; Department of Kinesiology and Health Education, University of Texas at Austin, Austin, TX, USA; Department of Mechanical and Industrial Engineering, Northeastern University, Boston, MA, USA; George W. Woodruff School of Mechanical Engineering, Georgia Institute of Technology, Atlanta, GA, USA; Insitute for Robotics and Intelligent Machines, Georgia Institute of Technology, Atlanta, GA, USA; School of Biological Sciences, Georgia Institute of Technology, Atlanta, GA, USA; Wallace H. Coulter Department of Biomedical Engineering, Emory University and Georgia Institute of Technology, Atlanta, GA, USA; Department of Rehabilitation Medicine, Division of Physical Therapy, Emory University, Atlanta, GA, USA

**Keywords:** exoskeleton control, balance augmentation, sensorimotor feedback

## Abstract

Wearable exoskeletons are a promising tool for augmenting balance and reducing fall risk. Recent work suggests that active ankle exoskeletons need to act faster than the human to improve reactive balance control. However, the magnitude of exoskeleton torque that is best for improving reactive balance remains unknown. Drawing from the optimal torque for minimizing metabolic expenditure, we hypothesized that reactive balance would improve with increased exoskeleton torque. Participants wearing bilateral ankle exoskeletons were instructed to maintain standing balance during 15cm backward support-surface perturbations. Three exoskeleton plantarflexion torque conditions were tested: NO (Off), LOW (15Nm), or HIGH (30Nm). LOW torque improved balance performance compared to NO torque (*p*<0.001), with a 7±3% decrease in peak center of mass (CoM) displacement. Although HIGH torque caused a 9±11% decrease in peak CoM displacement compared to NO torque (*p*=0.12), it was not significant due to high intersubject variability. Whereas LOW torque decreased peak CoM displacement in all (range: -0.2 to -1.6cm), HIGH torque only decreased it in some (range = 1.2 to -2.6cm). The change in CoM displacement from LOW to HIGH torque was associated with balance ability, quantified by the narrowing beam test (R^2^=0.29, p=0.06), while this relationship didn’t meet conventional statistical significance, likely due to the small sample size, it suggests that higher levels of exoskeleton torque may hinder balance performance in individuals with better balance ability. Taken together, more exoskeleton torque is not always better for balance, highlighting a potential need to personalize exoskeleton torque for balance augmentation.

## BACKGROUND

Humans are naturally unstable. Maintaining an upright, bipedal posture requires continuous muscle activity to keep the body’s relatively high center of mass (CoM) aligned over the base of support (Loram and Lakie, 2002; Morasso and Sanguineti, 2002; Winter, 1995). Thus, it is no surprise that falls, a failure to maintain balance, are a leading cause of traumatic injury and are exacerbated with aging or neuromuscular pathologies (Aschkenasy and Rothenhaus, 2006; Winter, 1995). Falls are not only costly to healthcare systems (Florence et al., 2018), but they also have psychosocial effects that impact independence and quality of life (O’Donnell et al., 2008). As such, there is a critical need to reduce the prevalence and burden of falls. Robotic exoskeletons have recently emerged as a promising solution to enhance balance and potentially reduce fall risk (Afschrift et al., 2023; Bayón et al., 2022; Emmens et al., 2018; Farkhatdinov et al., 2019; Monaco et al., 2017; Zhang et al., 2018). However, given the nascent objective of developing balance-augmenting exoskeletons, how exoskeleton control parameters should be determined, including the magnitude of exoskeleton torque, remains unclear. Here we evaluate how different magnitudes of bilateral ankle exoskeleton torque influence balance performance.

We do not know how the magnitude of applied exoskeleton torque affects balance performance. While limited work has focused on optimizing exoskeleton control parameters for balance augmentation, significant advances have been made in determining the optimal control parameters for minimizing metabolic cost during steady-state walking using active ankle exoskeletons (Ingraham et al., 2022; Poggensee and Collins, 2021; Quinlivan et al., 2017; Slade et al., 2022; Zhang et al., 2017). During walking, applying higher levels of exoskeleton torque— up to approximately 50 Nm (Slade et al 2022, Zhang et al 2017)—are optimal compared to lower levels of torque (Quinlivan et al., 2017). However, it is unclear whether a similar principle applies to balance augmentation, as the optimal torque timing differs between walking and balance. Namely, to minimize metabolic cost, exoskeleton torque at the hip or ankle should be delayed relative to the biological torque by approximately 100 ms (Molinaro et al., 2024; Young et al., 2017). In contrast, for balance augmentation, exoskeleton torque at the ankle should be applied faster than the physiological feedback response to enhance balance performance (Beck et al., 2023), at least in healthy young adults. Differences such as these indicate that we may be unable to extrapolate findings from other contexts when determining optimal exoskeleton control parameters for balance augmentation.

Another important consideration when determining the magnitude of exoskeleton torque for balance augmentation may be interindividual variability. For example, in walking, human-in-the-loop optimization for an ankle exoskeleton found interindividual differences in the optimal torque profile for energy optimization (Poggensee and Collins, 2021). Moreover, when tuning parameters based on user preference, significant interindividual variability exists in self-selected peak torque and timing during the gait cycle (Ingraham et al., 2022). Similarly, the optimal magnitude of exoskeleton torque may be influenced by task difficulty, someone’s inherent balance ability (e.g., an individual’s capacity to maintain balance)—which we quantify via the beam walking score (Sawers and Hafner, 2018)—or individual differences in the balance-correcting response. The same perturbation magnitude will result in a greater biomechanical challenge in shorter participants due to greater angular displacement of the CoM relative to the feet (Payne et al., 2019). Additionally, the balance correcting response arises from highly coordinated spinal and supraspinal sensorimotor feedback pathways (Boebinger et al., 2024; Jakubowski et al., 2025). Prior work has shown that the degree to which cortical brain activity is elicited is related to balance ability and task difficulty (Boebinger et al., 2026; Payne et al., 2019). Although direct measures of cortical activity are not widely available, recent work shows that cortical contributions to balance can be estimated based on muscle activity (Boebinger et al., 2024) and joint torque (Jakubowski et al., 2025). However, it is unclear if these balance-related factors are linked to performance within an exoskeleton.

In this study, we sought to evaluate how increasing exoskeleton torque magnitude affects reactive balance in healthy young adults. Drawing from prior research on ankle exoskeletons aimed at minimizing metabolic cost during walking (Slade et al., 2022; Zhang et al., 2017), and considering the torque limit of our device (Dephy ExoBoots with a peak torque of 30 Nm), we predicted that higher magnitudes (30 Nm) of applied ankle plantarflexion torque would improve balance compared to lower torque magnitudes (15 Nm). In addition, we aimed to examine whether interindividual variability in reactive balance, as quantified by peak CoM displacement, was related to participant height, balance ability, or cortical involvement during perturbations. Identifying whether such individual differences are related to the torque provided by an exoskeleton may help determine how to best personalize exoskeleton torque for balance augmentation.

## METHODS

### Participants

Ten healthy young adults (6 males and 4 females; age 26 ± 3 years; height 1.77 ± 0.11 m; mass 79 ± 14 kg) participated in this study after providing informed written consent in accordance with the Emory Institutional Review Board (IRB00082414).

### Ankle exoskeleton control

Participants wore bilateral ExoBoots (EB504; Dephy Inc., USA). The ExoBoots had a commercial boot with a carbon fiber plate embedded in the midsole and a laterally mounted quasi-direct drive actuator that could apply a maximum of 30 Nm of plantarflexion torque. The motor was connected to the ankle joint via a nonlinear, unidirectional cable-based transmission. Open-loop torque control was implemented via closed-loop current control, and manufacturer-specified motor constants were used to calculate the motor torque (characterized in (Lee et al., 2019)). Ankle torque was then determined from the motor torque and our internal characterization of the nonlinear transmission ratio (between 10:1 and 14:1 in the tested range of motion).

The ExoBoots were controlled via a Raspberry Pi 4B that reads in sensor data and commanded actuator torque at 200 Hz via USB (Raspberry Pi Foundation, UK). Custom Python scripts were used to control the ExoBoots. ExoBoots were commanded to provide plantarflexion torque after onboard accelerometers detected the onset of the backward support surface translation. The signals from onboard accelerometers were high-pass filtered with a second-order, 1 Hz Butterworth filter. The perturbation detection threshold was 147 cm/s^2^ in the posterior direction and less than 98 cm/s^2^ in both medial and lateral directions in either ExoBoot.

Three ExoBoot conditions were evaluated: NO torque (OFF), LOW torque (peak of 15 Nm of applied plantarflexion torque), and HIGH torque (peak of 30 Nm of applied plantarflexion torque) (Fig. 1). In the NO torque condition, 1 Nm of plantarflexion was applied to maintain cable engagement. In the LOW and HIGH torque conditions, a piecewise cubic hermite interpolating polynomial with torque defined as a function of time initiated immediately after perturbation detection (time vector: time= (0, 50, 200 ms) and torque vector: torque= (0, Peak Torque, 0 Nm)).

**Figure 1.**
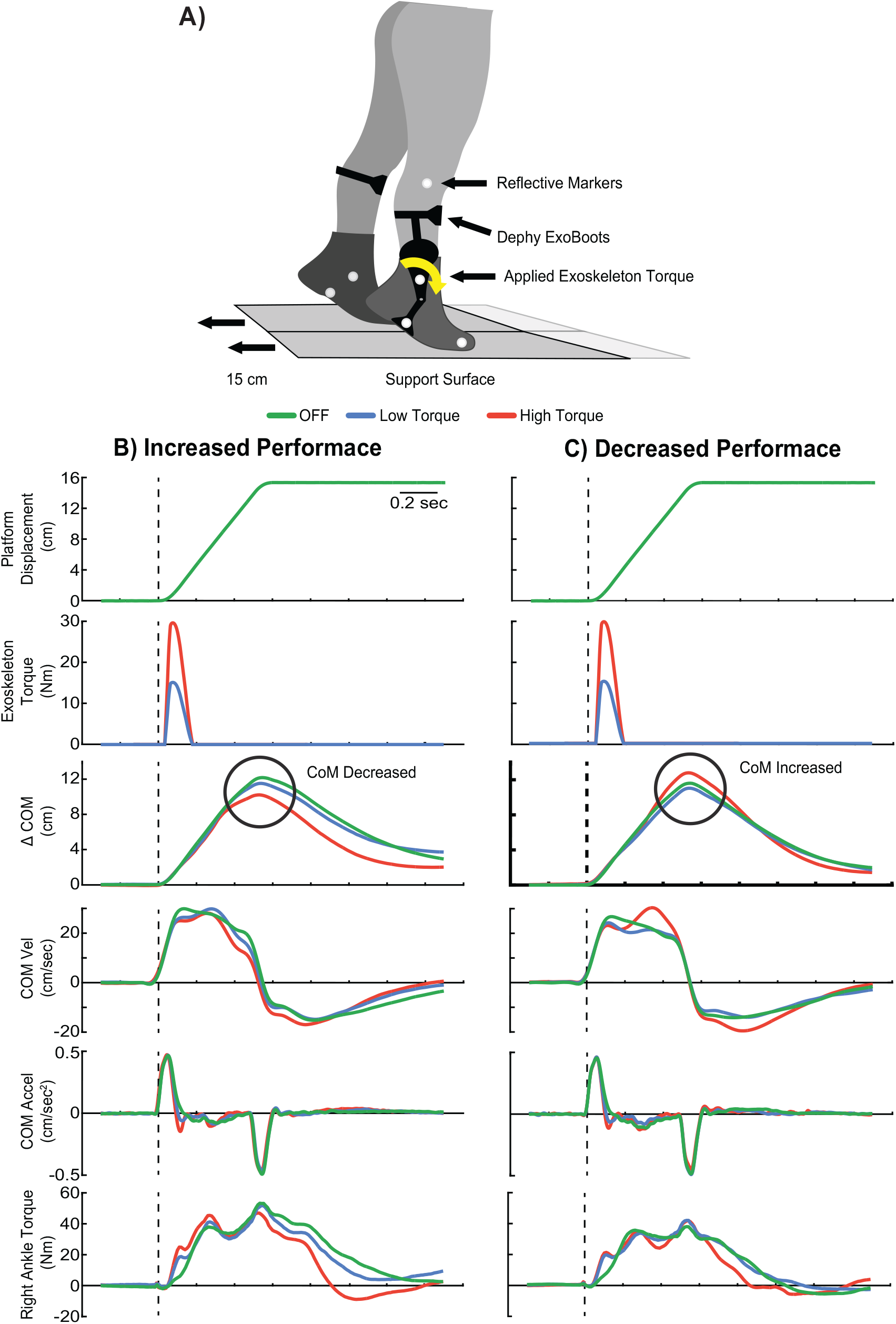
(A) A schematic of the experimental set-up. Participants maintained balance in response to a 15 cm backward support surface perturbation while wearing the Dephy ExoBoots (black), which applied either NO (green), LOW (15 Nm – blue), or HIGH (30 Nm - red) plantarflexion torque. (B) Representative data from one participant, where the center of mass (CoM) decreased during the HIGH torque (red) condition compared to the LOW (blue) torque condition, indicative of improved balance performance. (C) Representative data from one participant, where the center of mass (CoM) increased during the HIGH torque (red) condition compared to the LOW (blue) torque condition, indicative of worsened balance performance.

### Experimental set-up and Protocol

Participants were instructed to maintain standing balance in response to ramp-and-hold support surface translations. Participants stood on two independent force plates (AMTI, Watertown, MA, USA) embedded in a custom platform (Factory Automation Systems, Atlanta, GA) (Fig. 1). Ground reaction forces were collected at 1000 Hz. We placed 33 markers in accordance with a modified version of the Vicon Plug-in Gait model (Welch and Ting, 2008) that included additional foot markers (fifth metatarsal, medial and lateral heel, and medial malleolus).

The data presented here are part of a larger study where participants experienced a total of 106 perturbations (Beck et al., 2023). Participants completed 4 backward ramp-and-hold perturbations at 15 cm in each ExoBoot condition (NO, LOW, and HIGH) in a randomized fashion. To mitigate adaptation and anticipation, participants experienced 8 cm catch trials (e.g., forward perturbations) randomly interspersed within the perturbation set. A 5-minute seated rest break followed every 20 perturbations to mitigate fatigue. Perturbation trials that elicited a stepping response or trials where participants uncrossed their arms were considered invalid. Stepping responses were identified as trials in which the magnitude of ground reaction forces for either leg dropped below 10 N.

### Data processing

Processing and analysis were performed using custom-written software in MATLAB (Mathworks, Natick, MA, USA). All marker data and ground reaction forces from both force plates were used to estimate joint kinematics and kinetics. All marker data were filtered using a fourth-order low-pass filter with a 10 Hz cutoff, while ground reaction forces were filtered using a fourth-order low-pass filter with a 50 Hz cutoff. Inertial artifacts that arose from translating the platform were removed (Hnat et al., 2018; Preuss and Fung, 2004). Joint kinematics and kinetics were estimated using the inverse kinematics and inverse dynamics toolboxes in OpenSim (Gait 2892 model), respectively (Delp et al., 2007). The CoM acceleration was calculated as the ground reaction forces divided by the participant’s mass. CoM displacement and velocity were calculated as the weighted sum of all segmental masses as previously done (McKay et al., 2021; Safavynia and Ting, 2013; Welch and Ting, 2014; Welch and Ting, 2009). CoM displacement, velocity, and acceleration were all taken relative to the movement of the platform. CoM displacement and velocity were upsampled using linear interpolation to 1000 Hz for all further analysis. Peak CoM displacement was estimated as the max of the CoM displacement. (Fig 1).

### Narrowing Balance Beam Walking Test

Balance ability was assessed using a narrowing-beam walking task (Sawers and Hafner, 2018) that was modified to be more challenging to account for the relatively higher balance ability of healthy young adults. The task uses three sequential beam segments (one 4 cm wide and two 2 cm wide), each 182.9 cm long. Participants completed five trials of walking along a progressively narrowing beam with their arms crossed across their trunk. Trials were terminated when participants stepped off the beam or uncrossed their arms. Distance traveled was recorded for each trial, and scores from trials 3–5 were normalized by the total beam length (548.7 cm) and averaged to obtain a final balance ability score (Sawers and Hafner, 2018).

### Delayed center of mass (CoM) feedback model

To evaluate interindividual variability in the level of cortical involvement during the balance-correcting response, we used a previously developed delayed CoM feedback model that reconstructs the balance-correcting ankle torque response and can characterize different feedforward and feedback contributions to the balance-correcting response (Jakubowski et al., 2025). The model reconstructs joint torque based on CoM kinematics, where CoM kinematics (displacement, velocity, and acceleration) are multiplied by feedback gains (*K*_*D*_, *K*_*V*,_ *K*_*A*_) and a common delay (*λ*) (Supplemental Figure 1). We have previously demonstrated that two feedback loops are required to capture all salient features of the ankle torque response (Jakubowski et al., 2025). We tuned the gains and delays within each loop to optimize the fit between the inverse dynamics-derived and the CoM feedback-derived ankle torques for each participant during the no torque condition (Supplemental Figure. 2). We note that the delayed CoM feedback model could not be used when the exoskeleton was applying plantarflexion torque (LOW and HIGH torque conditions) because the biological ankle and applied exoskeleton torque could not be dissociated. All optimizations were performed in MATLAB R2023b.

### Statistical analysis

We evaluated how exoskeleton torque magnitude (NO, LOW, or HIGH) affected peak CoM displacement, our measure of balance performance. We compared peak CoM displacement using a linear mixed-effects model, with subject treated as a random factor and exoskeleton condition as a fixed factor. We used a restricted maximum-likelihood method and Satterthwaite corrections to account for the small sample size and reduce Type 1 errors (Luke, 2017).

We evaluated whether height, balance ability (beam-walking score), and optimized gains from the delayed CoM feedback model were related to the observed interindividual variability during the HIGH torque condition. To do this, we calculated the difference in peak CoM displacement between LOW and HIGH torque (*ΔCoM*) and evaluated whether that difference correlated with height, balance ability, and the CoM feedback model gains via a linear regression model. All statistical analyses were performed in MATLAB R2023b. Significance was set a priori at α=0.05. Bonferroni corrections were performed for all post-hoc analyses, with Bonferroni-corrected p-values reported. All metrics were reported as mean ± standard deviation.

## RESULTS

### More exoskeleton torque improved balance performance in some but not all participants

While the lower magnitude of exoskeleton torque improved standing balance performance across all individuals, the higher torque magnitude only provided additional benefits for roughly half the participants (Fig. 2). The LOW torque condition significantly improved balance performance (*p* < 0.001), with peak CoM displacement being 7±3% lower during the LOW torque (10.3±0.9 cm) compared to the NO torque condition (11.1±0.8 cm). However, there was only a 2±10% decrease in peak CoM displacement in the HIGH torque condition (10.1±1.5 cm) compared to the LOW torque condition (p=1). Although the peak CoM displacement in the HIGH torque condition was 9±11% lower than the NO torque condition, this difference was not statistically significant (p=0.12). These results reflect intersubject differences in the response to the different exoskeleton conditions. While all participants decreased peak CoM displacement in the LOW vs NO torque conditions (range: -0.2 to -1.6 cm), in the HIGH vs LOW torque conditions participants showed either a further decrease, an increase, or relatively no change in peak CoM displacement (range: 1.2 to -2.6 cm).

**Figure 2.**
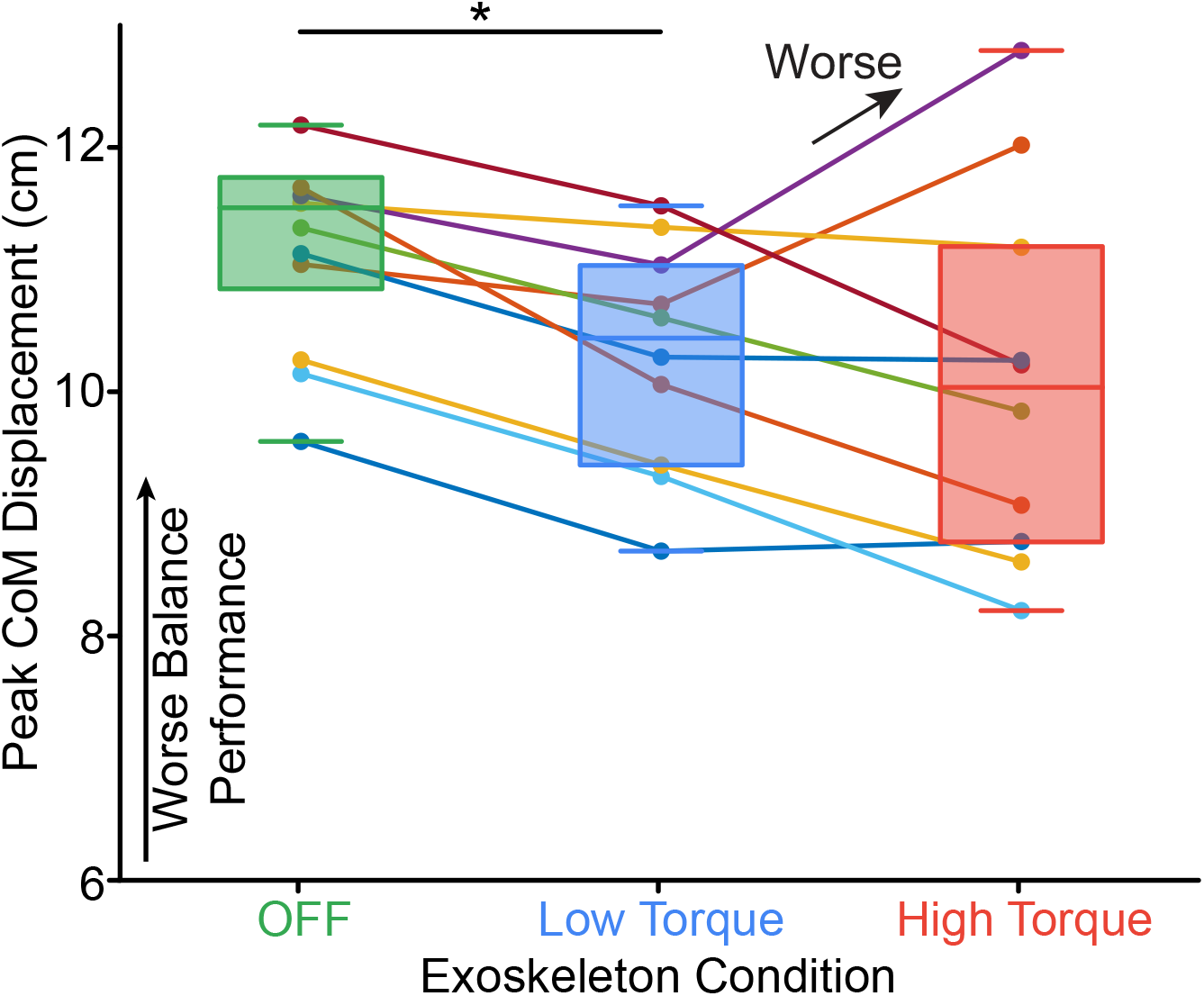
For all participants, the LOW torque significantly decreased peak CoM displacement; however, there was high interindividual variability in the response to the HIGH torque, and thus, no significant changes compared to either NO (green) or LOW (blue) torque. The colored boxes represent the median (middle line) and the 25th and 75th percentiles (bottom and top edges, respectively) for each exoskeleton condition. The colored dots and lines represent each participant. The asterisks indicate a significant difference (p < 0.05).

### Factors contributing to high interindividual variability

We found a trend in which individuals with higher balance beam scores showed decreased balance performance between LOW and HIGH exoskeleton torque (R^2^=0.29, p=0.06). This nearly significant finding was based on correlating narrowing beam walking scores (average normalized score=0.2 ± 0.2) (Fig. 3A) with ΔCoM, the difference in peak CoM displacement between LOW and HIGH torque.

**Figure 3.**
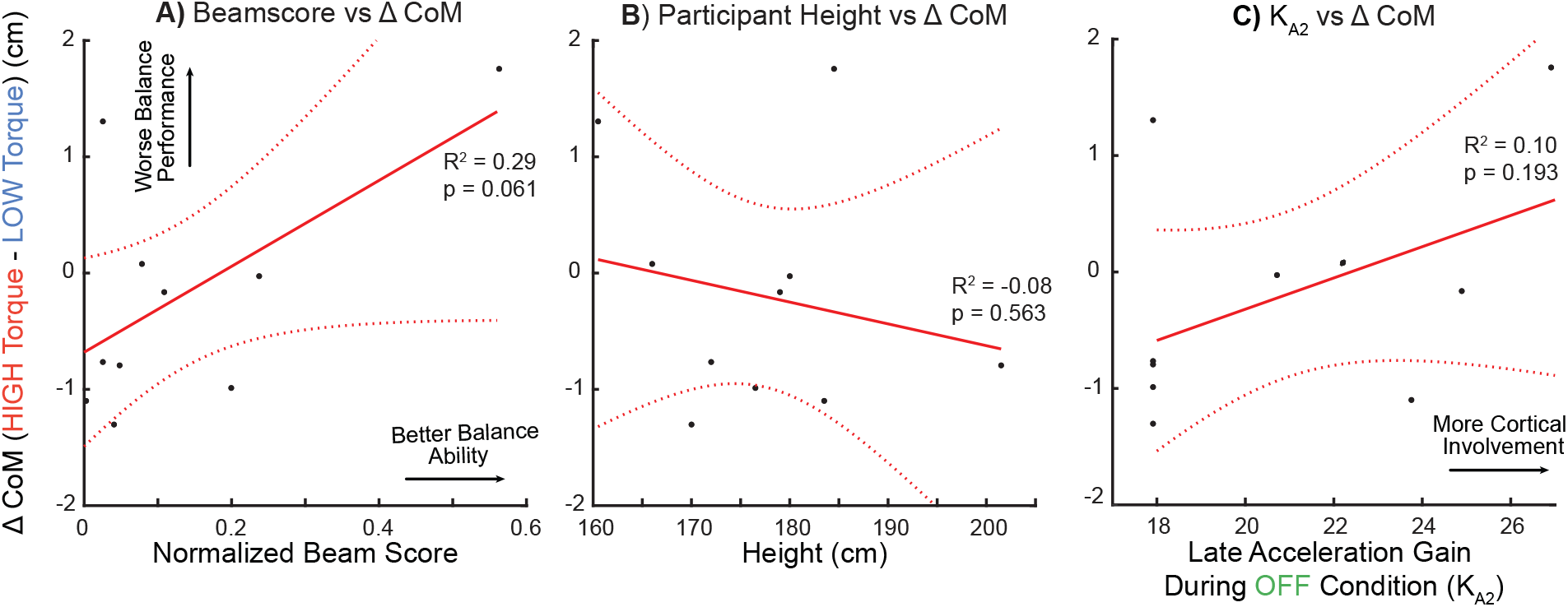
(A) There was a trend between the difference in CoM displacement between the HIGH torque and LOW torque conditions (Δ*CoM*) and the scores of the narrowing balance beam walking test, indicating that individuals with higher balance ability performed worse during the HIGH exoskeleton torque condition. (B) ΔCoM was not related to participant height. (C) ΔCoM was not related to the optimized acceleration gain from the late feedback loop (KA2), which estimates the level of cortical involvement during the balance correcting response. (A-C) the solid red line indicates the linear fit, with the region outlined by the dotted red lines being the 95% confidence intervals. The black dots represent each participant.

In contrast, there was no relationship between subject height and ΔCoM (R^2^=-0.08 and p=0.56) (Fig. 3B).

Additionally, there was no relationship between the cortical contributions to balance and ΔCoM. The delayed CoM feedback model accurately reconstructed the ankle torque response during the NO torque condition and characterized the specific feedback pathways generating the overall torque response, as previously demonstrated (Jakubowski et al., 2025). Consistent with prior findings, the balance-correcting ankle torque response consisted of only feedback contributions, one “early” feedback loop (average *λ*_*1*_=107 ± 15 ms; e.g., subcortical response), and one “late” feedback loop (average *λ*_*2*_=196 ± 33 ms; e.g., cortical response). *K*_*A2*_, the feedback gain within the “late” feedback loop, which is thought to represent the level of cortical involvement, was not related to Δ*CoM* (*R*^*2*^=0.10 and *p*=0.19) (Fig. 3C), nor were other gains from the CoM feedback model (all p>0.17, supplemental material).

## DISCUSSION

We found that balance performance was not consistently improved across individuals as exoskeleton torque was increased. Although all participants improved balance at a lower level of torque, doubling that torque—the maximum allowed by our device—did not always improve balance performance (5 improved, 2 worsened, 3 stayed the same). Surprisingly, our data suggest that individuals with better balance ability were less likely to benefit from higher exoskeleton torque, indicating that higher exoskeleton torque may hinder individuals with better balance. Collectively, our work highlights the need for personalized exoskeleton controllers that account for individual variability and the need for further investigation into factors that may determine optimal control parameters for balance augmentation.

### More exoskeleton torque is not always better

There may be differences in control policies between using exoskeletons for reducing metabolic cost during steady-state walking and for balance augmentation. While the lower magnitude of exoskeleton torque improved standing balance performance across all individuals, the higher torque magnitude only provided additional benefits for half our participants (Fig. 2).

This finding contradicts our central hypothesis and findings from studies using exoskeletons for energy optimization during steady-state walking. During walking, the optimal peak torque is approximately 0.7 Nm/kg, corresponding to approximately 50% of peak biological torque (Slade et al., 2022; Zhang et al., 2017). In reactive balance, 30Nm corresponds to approximately 0.4 Nm/kg of torque and about 65% of the peak reactive ankle torque. Thus, while we applied a lower torque percentage scaled by body weight, it represented a larger portion of the biological torque. It is unclear if torque should be scaled to biological torque for each task or by body weight. Combined with prior findings (Beck et al., 2023), our results suggest that there may be differences in control parameters between the two applications; future work is warranted to determine which, if any, policies transfer between applications.

Our data support the idea that interindividual differences are critical when optimizing exoskeleton torque for balance. Prior studies that used human-in-the-loop optimization to optimize the torque profiles that best minimized energy expenditure during steady-state walking found interindividual differences in the peak torque (Poggensee and Collins, 2021). Similarly, we found that the same peak torque magnitude could assist balance in some individuals, while worsening balance in others. As worsening balance might even increase the risk of falls, the importance of poorly optimized exoskeleton torque may have even greater significance than in walking. Thus, techniques to optimize exoskeleton torque, as has been done in studies that sought to minimize metabolic cost during walking (Slade et al., 2022; Zhang et al., 2017), will be critical to developing a deployable balance-augmenting exoskeleton.

Perhaps surprisingly, our work suggests that torque provided by the exoskeleton may hinder reactive balance performance in those with better balance ability. Balance correction relies on highly coordinated sensorimotor feedback pathways that integrate sensory information at both the spinal and supraspinal levels (Boebinger et al., 2024; Jakubowski et al., 2024). The balance correcting response is continuously refined through experience and adapted to an individual’s biomechanics and motor coordination patterns (Torres-Oviedo and Ting, 2007). Individuals with better balance ability may possess highly adapted balance-correcting responses that efficiently generate the precise timing and magnitude of corrective torques required to maintain balance. Further, recent work suggests that individuals with better balance ability are more sensitive to balance perturbations (Kerr et al., 2026). In these individuals, the exoskeleton could have been an additional “disturbance”, interfering with their intrinsic balance-correcting responses.

Future work should also investigate the role of motor learning in determining exoskeleton torque magnitude for balance augmentation. Prior work has found improvement in standing balance performance with higher exoskeleton torque magnitudes when the torque was consistently applied across trials (Eveld et al., 2024). In contrast, our experimental paradigm minimized opportunities for adaptation by randomizing exoskeleton torque conditions throughout the experiment. Prior studies that aimed to minimize metabolic cost during steady-state walking have found a significant decrease in metabolic cost after training, but there was interindividual variability in the length of training required (Poggensee and Collins, 2021). Similarly, learning to effectively use exoskeletons for balance augmentation may be required, as well as consideration of interindividual differences in their response to training. Training time required may be determined by how much individuals initially “resist” the exoskeleton (Gordon and Ferris, 2007) and by demographic factors such as age and the presence of neuromuscular disease, both of which have been linked to decreased motor learning capacities (King et al., 2013).

### Limitations

Our study has several potential limitations. First, the time-synchronized applied exoskeleton torque was not measured during the LOW and HIGH torque conditions. Therefore, since the biological torque and the applied exoskeleton torque could not be dissociated, we could not apply the CoM feedback model during the LOW and HIGH torque conditions to estimate cortical involvement during those conditions. Future work should investigate whether the cortical gain changes with the magnitude of applied exoskeleton torque. Additionally, the ankle exoskeletons only applied a plantarflexion torque to the user. Thus, we only evaluated the balance-correcting response during backward support-surface perturbations that require a corrective plantarflexion torque. Similarly, only a single perturbation magnitude was evaluated (15 cm). Future work should incorporate perturbations of various magnitudes and directions with an exoskeleton capable of applying both plantarflexion and dorsiflexion torque to determine the optimal exoskeleton control parameters and the generalizability of those parameters. Finally, the analyses relating ΔCoM to height, balance ability, and feedback gains were performed in a small sample and may be underpowered. These findings should therefore be interpreted as hypothesis-generating.

## CONCLUSION

A higher magnitude of exoskeleton torque is not always better for balance performance. Some individuals benefit more from using lower magnitudes of exoskeleton torque, while others benefit from higher magnitudes of exoskeleton torque for balance performance. Those with better balance may in fact be hindered by too much exoskeleton torque. While future investigation into what factors drive performance in the exoskeleton is required, our results highlight that personalized exoskeleton controllers may be needed when augmenting balance performance.

## DATA AVAILABILITY

The data from the current study are available from the corresponding author upon reasonable request.

## GRANTS

This publication was supported by grant number 2127509 from the NSF and American Society for Engineering Education, National Institutes of Health grants F32 HD112128, F32 AG063460, R01 HD046922, R01 HD090642, and McCamish Parkinson’s Disease Innovation Program. Its contents are solely the responsibility of the authors and do not necessarily represent the official views of the National Science Foundation, American Society for Engineering Education, National Institutes of Health, or McCamish Foundation.

## DISCLOSURES

The authors declare no conflicts of interest, financial or otherwise.

### LIST OF ABBREVIATIONS

Abbreviation: Definition
CoM: Center of Mass
RMSE: Root Mean Square Error

## SUPPLEMENTAL

### Extended methods - Delayed CoM feedback model

To use the delayed CoM feedback model, all trials from the NO torque condition were averaged to generate the average CoM kinematics and inverse dynamics-derived ankle joint torque. The average curves were then used for all subsequent analyses. To isolate reactive ankle torque, background torque (defined as the mean ankle torque one-second before platform movement) was subtracted before optimizing gains and delays. For each of the two loops, optimization using *fmincon*.*m* was performed to identify *K*_*Di*_, *K*_*Vi*,_ *K*_*Ai*_, and loop to prevent the λ_i_, where *i* indicates the *i* th loop. Parameter bounds were imposed on each loop to prevent the algorithm from searching outside a physiologically relevant space and reconstructing the same ankle torque curve features. During the fitting process, the fit of each loop was evaluated. If the loop poorly reconstructed the data, hand-tuning of the bounds was performed to achieve the best model as quantified by the coefficient of determination (*R*^***2***^) and variance accounted for (*VAF*). After two separate optimizations identified the best gains for each loop, the gains and delays were concatenated as the initial guess for a final optimization. The final optimization concurrently optimized the gains for both loops. In this optimization, the lower and upper bounds were restricted to ±10% of the initial optimization, while the delays were restricted to ±10ms of the initial optimization.

**Supplementary Figure 1.**
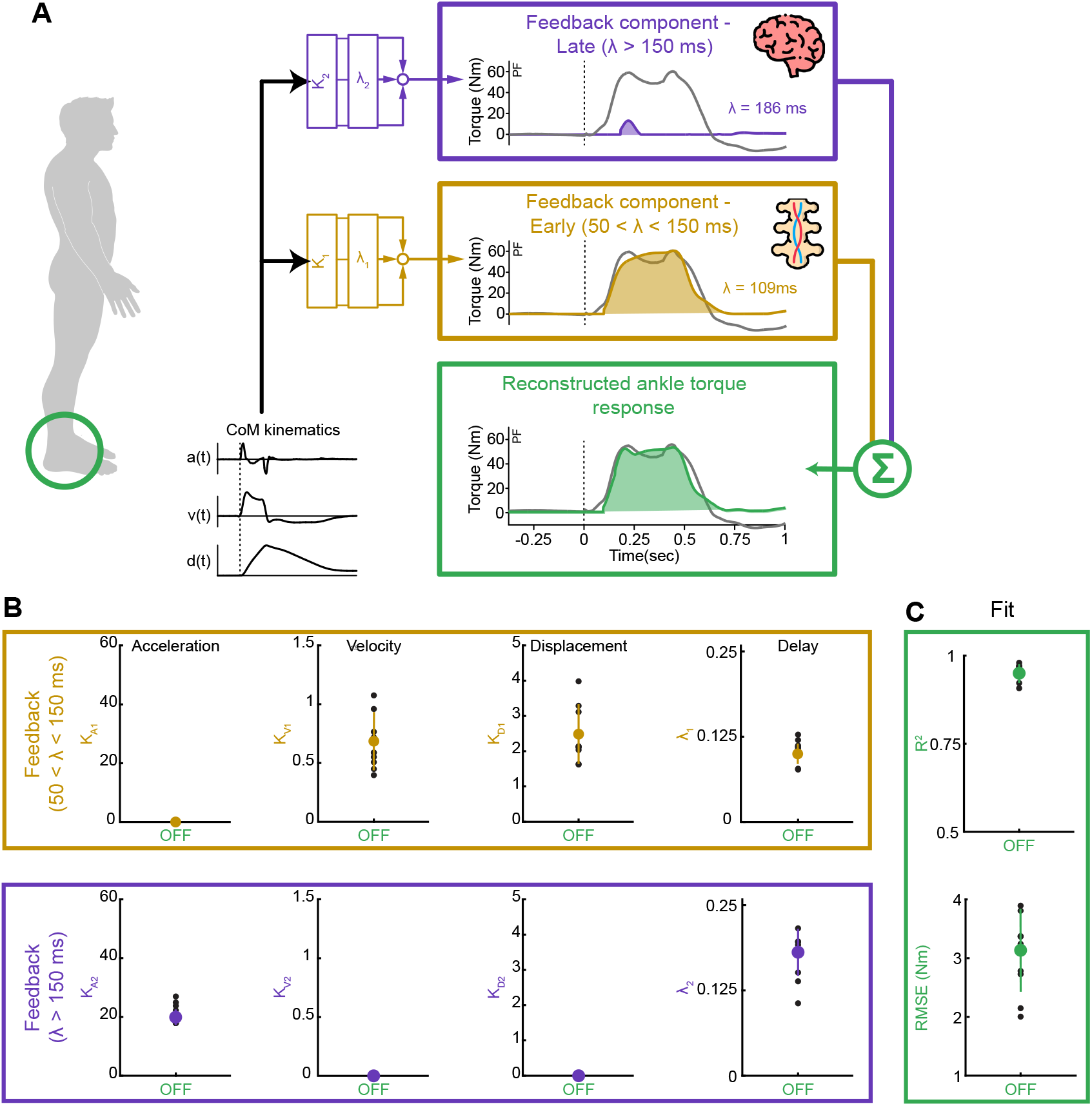
(A) The delayed center of mass (CoM) feedback model for the ankle torque response, indicating that the response is mediated by an “early” (delay < 150 ms; gold) and “late” (delay > 150 ms; purple) feedback component. These two components are summed together, resulting in the overall ankle torque response (green). (B) The delayed CoM feedback model gains for each loop. *K*_*A*_, *K*_*V*_, and *K*_*D*_ represent the CoM displacement, velocity, and acceleration gains, while λ is the time delay. (C) The delayed CoM feedback model reconstructed the torque response well, with a high *R*^*2*^ and low *RMSE*. (B-C) The colored dots represent the mean and the colored vertical lines represent the standard deviation of the optimized gains. The black dots represent each participant.

**Supplementary Figure 2.**
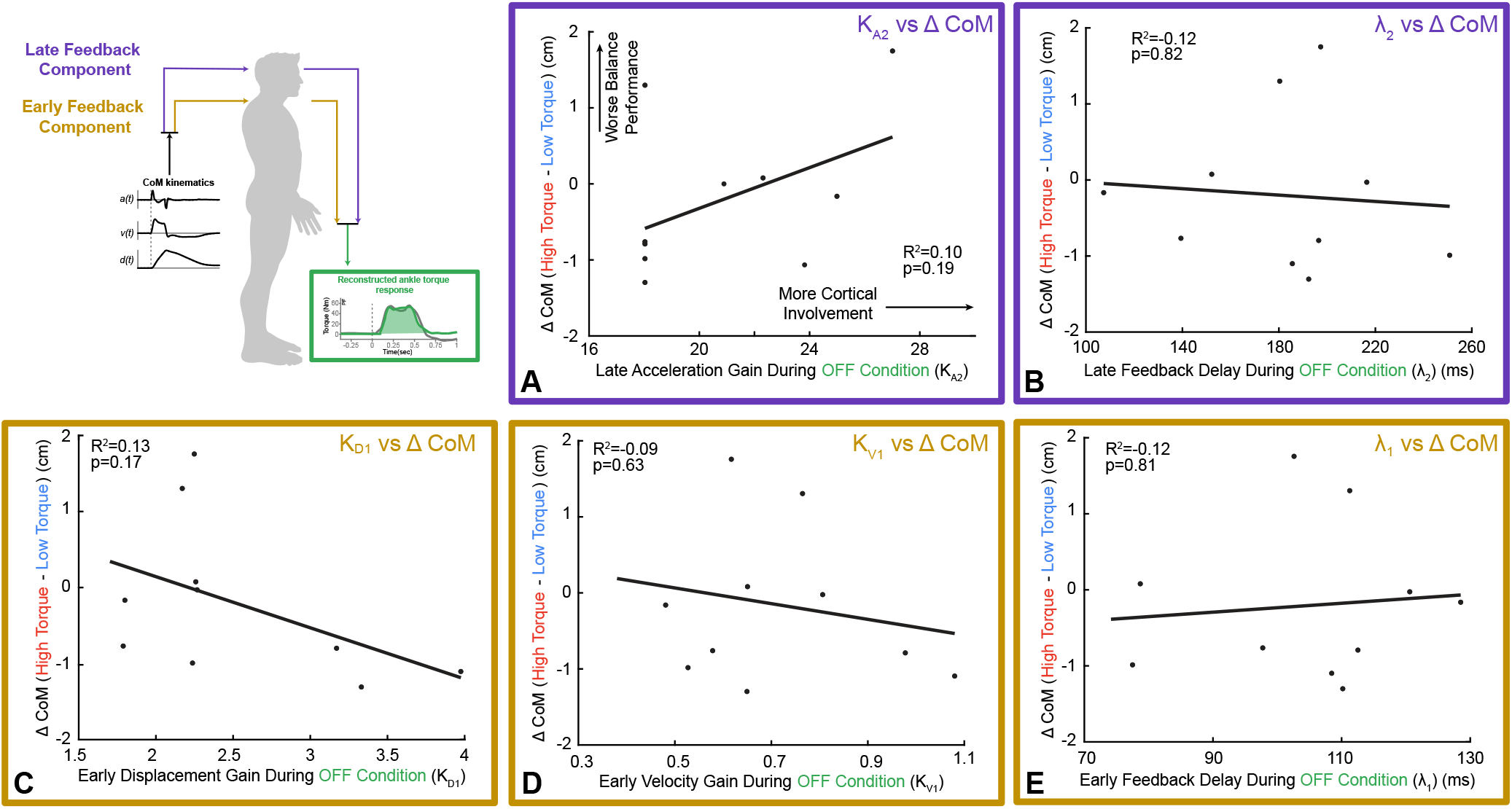
(A-E) The difference in CoM displacement between the HIGH torque and LOW torque conditions (Δ*CoM*) was not related to any of the optimized feedback gains or delays. The solid black line indicates the linear fit. The black dots represent each participant.

## Notes

### Competing Interest Statement

The authors have declared no competing interest.

